# Illustrating phylogenetic placement of fossils using RoguePlots: An example from ichneumonid parasitoid wasps (Hymenoptera, Ichneumonidae) and an extensive morphological matrix

**DOI:** 10.1101/425090

**Authors:** Seraina Klopfstein, Tamara Spasojevic

## Abstract

The fossil record constitutes the primary source of information about the evolutionary history of extant and extinct groups, and many analyses of macroevolution rely on fossils that are accurately placed within phylogenies. To avoid misinterpretation of the fossil record, especially by non-palaeontologists, the proper assessment and communication of uncertainty in fossil placement is crucial. We here use Bayesian morphological phylogenetics to evaluate the classifications of fossil parasitoid wasps (Hymenoptera, Ichneumonidae) and introduce ‘RoguePlots’ to illustrate placement uncertainty on the phylogeny of extant taxa. Based on an extensive, newly constructed morphological matrix of 222 characters in 24 fossil and 103 extant taxa, we test three different aspects of models of morphological evolution. We find that a model that includes ordered characters, among-character rate variation, and a state-space restricted to observed states achieves the highest marginal likelihoods. The individual RoguePlots reveal large differences in confidence in the placement of the different fossils and allow some refinements to their classification: *Polyhelictes bipolarus* and *Ichninsum appendicrassum* are moved from an uncertain subfamily placement to Pimplinae, *Plectiscidea lanhami* is transferred to *Allomacrus* in Cylloceriinae (*Allomacrus lanhami*, comb. nov.), *Lithotorus cressoni* is moved from Diplazontinae to Orthocentrinae, and we note uncertainty in the generic placement of *Xanthopimpla*? *messelensis*. We discuss potential artefacts that might result in biased posterior probabilities in Bayesian morphological phylogenetic analyses, pertaining to character and taxon sampling, fossilization biases, and model misspecification. Finally, we suggest future directions both in ichneumonid palaeontology, in the modelling of morphological evolution, and in the way Bayesian phylogenetics can improve both assessment and representation of fossil placement uncertainty.

## Introduction

The fossil record provides crucial information about the evolutionary history of a group and allows us, by facilitating the inference of absolute ages, to put said history into a palaeogeographic and palaeoecological context. Many analyses that use the fossil record rely on a firm placement of fossils in the phylogenetic or at least taxonomic context of extant species. This is especially true for the dating of molecular trees through fossil calibrations [1], but also for the inference of ancient interactions between different groups of organism [2,3] and for the study of evolutionary trends in morphological characters over large timescales [e.g., 4,5]. Unfortunately, the correct taxonomic interpretation of fossils can be a formidable challenge due to incomplete preservation, difficulties in the taphonomic interpretation, and long gaps in the fossil record [6,7,8]. The proper communication of this uncertainty is crucial in order to avoid future misinterpretations and overconfidence in analyses that are relying on reliably placed fossils.

Even though the “open nomenclature” framework [9] provides some means to express uncertainty in the placement of a fossil by way of adding a question mark after the genus name or explicitly not placing it in higher ranks (as ‘incertae familiae’, ‘incerti ordinis’, etc.), these tools are not very flexible and have been used inconsistently by different authors. Furthermore, the need for a binomial when describing new species often leads to new fossils being placed in extant genera without sufficient evidence for such placement. The justification for fossil classifications is often limited to brief mentions of character evidence, which might or might not be sufficient, depending on the group in question and preservation of the fossil. An alternative way to arrive at a well-grounded classification of fossils is to make use of a morphological phylogenetic analysis, which allows obtaining both a ‘best estimate’ for the position of the fossil and, even more importantly, some measure of support for said placement.

Until recently, the phylogenetic analysis of fossil taxa has been relying largely on parsimony as an analysis approach, while model-based approaches became the gold standard for molecular characters already quite a while ago [10,11]. Recent developments in the field of modelling the evolution of discrete morphological characters [e.g., 12,13,14,15] have readied morphological phylogenetics for stochastic frameworks such as maximum likelihood and Bayesian inference, a realization that has also impacted several palaeontological studies [16,17,18,19,20,21]. The stochastic framework allows for the explicit testing of alternative fossil placements, thus providing a vital tool for the assessment of its uncertainty. Bayesian approaches are especially interesting in this respect, as they directly provide posterior probabilities of a fossil attaching to different branches in a tree and thus allow assessing alternative placements in an intuitive way [11].

Due to the often severely incomplete preservation of fossils, uncertainty in their placement is typically very high. In a phylogenetic analysis, fossils thus often behave as ‘rogue’ taxa [22], i.e., taxa that drift around in the tree and thus substantially deteriorate the resolution and clade support values in a consensus tree [e.g., 23]. This is the case irrespective of whether a strict or majority-rule consensus is constructed, or whether it is based on a set of most parsimonious trees, bootstrap trees, or trees sampled during a Bayesian Markov Chain Monte Carlo (MCMC) analysis. To illustrate the phylogenetic placement of such ‘rogues’, they are best excluded before a consensus tree is created, which then might be much better resolved, and information of fossil attachment is then summarized on this tree. We here introduce a graphic representation of fossil placement uncertainty that summarizes Bayesian posterior probabilities, so-called ‘RoguePlots’, and apply it to assess the placement of 24 fossils in a group with a severely understudied fossil record: ichneumonid parasitoid wasps.

The Ichneumonidae are the largest family of parasitoid wasps and contain more than 25,000 described [24] and many more undescribed species [25]. Studying only the literature, one would come to the conclusion that the fossil record of the group is rather poor, as less than 300 species have been described to date [26]; however, this low number rather reflects the lack of palaeontologist working on that group, as ichneumonids are often among the more abundant insect groups found, at least at various Cenozoic localities [27,28,29]. Several recent studies have greatly furthered our understanding of the ichneumonid fossil record [7,28,30,31,32,33,34,35,36,37], but our knowledge of the evolutionary history of the group remains very patchy. Placement of fossil ichneumonids, even just into subfamilies, is often particularly difficult because of pronounced homoplasy in this group, which is probably the result of unrelated lineages attacking similar hosts in similar ecological niches [38]. Phylogenetic approaches have the potential to aid in identifying such homoplasies and thus improve fossil placement – or at least provide a realistic measure of their uncertainty. However, to date, no study has used a phylogenetic framework to place ichneumonid fossils in relation to their extant counterparts.

We here review 24 fossil ichneumonid species that we have described or revised recently [7,28,37], and formally test their phylogenetic placement. To this end, we assembled a large morphological matrix that includes both extant and fossil ichneumonid taxa, and perform Bayesian phylogenetic analyses under a range of evolutionary models. We use ‘RoguePlots’ to summarize the posterior distribution of a fossil’s position on the consensus tree of extant taxa, detail the implications of these results for their classification, and discuss the advantages of the Bayesian approach to fossil placement, especially with respect to the representation of phylogenetic uncertainty.

## Materials and Methods

### Taxon sampling and morphological matrix

We included 24 fossils that we have recently described or revised (Table 1) [7,28,37], with the taxonomic placements at the time supported by character evidence only. For most species, we studied the holotype and sometimes paratype directly, but in a few cases [listed in 7], we only obtained high-resolution photographs of the holotypes. No permits were required for the described study, which complied with all relevant regulations.

**Table 1.** Fossil taxa examined. The taxonomy reflects the status before our study; taxa marked with an asterisk before the genus name are transferred in this study.

| **Subfamily** | **Genus** | **Species** | **Epoch** | **Locality** |
| --- | --- | --- | --- | --- |
| Acaenitinae | *Mesoclistus?* | *yamataroti* | Early Eocene | Green River |
| Banchinae | *Glypta?* | *transversalis* | Early Eocene | Green River |
| Labeninae | *Trigonator* | *macrocheirus* | Early/Mid Eocene | Messel |
| Orthocentrinae | *\*Plectiscidea* | *lanhami* | Early Eocene | Green River |
| Phygadeuontinae | *Phygadeuon?* | *petrifactellus* | Early Eocene | Green River |
| Pimplinae | *Dolichomitus?* | *saxeus* | Early Eocene | Tadushi |
| Pimplinae | *Scambus* | *fossilobus* | Early/Mid Eocene | Messel |
| Pimplinae | *Scambus?* | *mandibularis* | Early Eocene | Green River |
| Pimplinae | *Scambus?* | *parachuti* | Early Eocene | Green River |
| Pimplinae | *Pimpla?* | *eocenica* | Early Eocene | Green River |
| Pimplinae | *Xanthopimpla* | *messelensis* | Early/Mid Eocene | Messel |
| Pimplinae | *Xanthopimpla* | *praeclara* | Early/Mid Eocene | Messel |
| Pimplinae | *Crusopimpla* | *tadushiensis* | Early Eocene | Tadushi |
| Pimplinae | *Crusopimpla* | *rediviva* | Late Eocene | Tadushi |
| Rhyssinae | *Rhyssella* | *vera* | Early/Mid Eocene | Messel |
| Tryphoninae | *Eclytus?* | *lunatus* | Early Eocene | Green River |
| incertae subfamiliae | *Carinibus* | *molestus* | Early Eocene | Green River |
| incertae subfamiliae | *Eopimpla* | *grandis* | Early Eocene | Green River |
| incertae subfamiliae | *\*Ichninsum* | *appendicrassum* | Early Eocene | Green River |
| incertae subfamiliae | *\*Lithotorus* | *cressoni* | Early Eocene | Green River |
| incertae subfamiliae | *Mesornatus* | *markovici* | Early/Mid Eocene | Messel |
| incertae subfamiliae | *\*Polyhelictes* | *bipolarus* | Early/Mid Eocene | Messel |
| incertae subfamiliae | *Tilgidopsis* | *haesitans* | Early Eocene | Green River |
| incertae subfamiliae | *Trymectus* | *amasidis* | Early Eocene | Green River |

Morphology of all fossils was studied in detail and characters were coded into a morphological matrix. Morphological nomenclature follows Townes [25], except for wing venation characters, which are based on Kopylov [36] with few modifications (Fig. 1).

**Figure 1.**
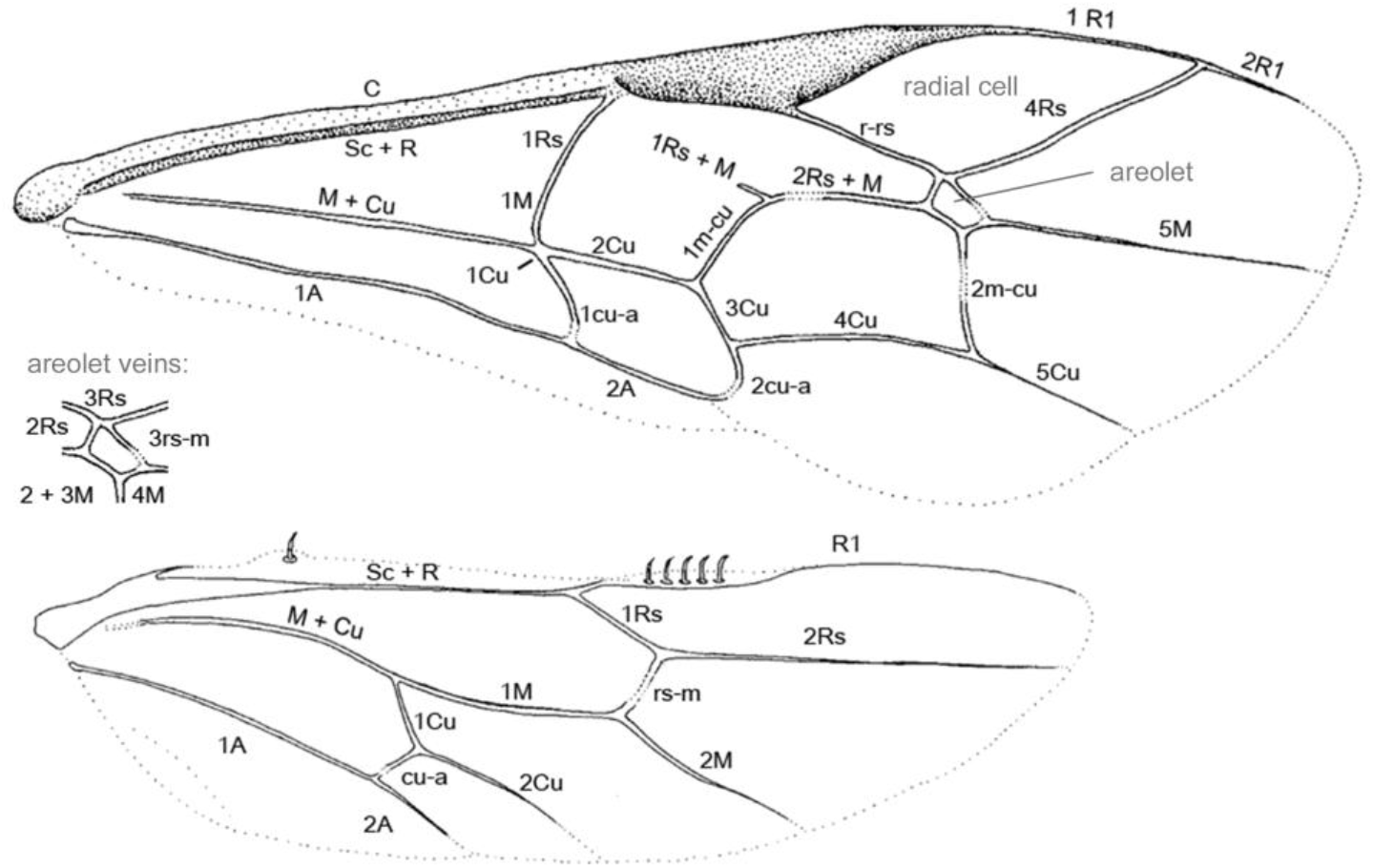
Wing vein and forewing cell nomenclature used in the current study.

The morphological matrix consists of 222 morphological characters of the adult insects scored for 103 extant and 24 fossil taxa. To this day, it represents the largest matrix of this kind composed for ichneumonids (Supplementary File S1). We aimed to get a good representation of the ichneumonid subfamilies (24 out of 42) while focusing on Pimpliformes, where we included 80 extant genera. In cases of heterogeneous and potentially non-monophyletic genera, we included representatives of several species groups. The full list of included taxa with specimen numbers and collection details is given in Supplementary File S2. Our matrix is partly based on the morphological matrix from Gauld et al. [39] (~50% of the characters), which had been designed to capture the morphological diversity within a single ichneumonid subfamily, the Pimplinae. It was complemented with characters from Bennett et al. (unpublished) (~15% of the characters), which had a broad but sparse taxon sampling across ichneumonids. In order to cover most of the morphological diversity of the taxa scored in this study, we have re-defined and expanded many of the character definitions, and added specific characters for previously underrepresented subfamilies. To maximise the morphological information coming from fossils, we added some characters that are not traditionally used in ichneumonid taxonomy but are often well (and sometimes exclusively) preserved in fossils, such as wing venation characters. The full list of characters and states including detailed descriptions is given in Supplementary File S3.

Out of the 222 morphological characters, a majority are multistate characters, while 34% are binary. The 13 continuous characters represent ratios of length measurements designed to capture the shape of structures (except of the fore wing length which was included as a measure of size). In order to be able to analyse them along with the remaining characters we eventually transformed them to ordered six-state discrete characters following the gap weighting approach as defined in Thiele [40]. Six states is currently the upper limit for ordered characters in the used phylogenetic software (see below). We took special considerations when scoring fossils (but also extant taxa) and coded uncertain interpretations as polymorphisms; initially, many questionable coding were noted with “?” (e.g., “1?”), and preliminary analyses with either keeping the states (“liberal approach”) or keeping the question marks (“conservative”) resulted in better support values when the liberal approach was applied, while not creating any supported conflict in tree topology compared to the conservative approach. We thus decided to continue with the more liberal interpretation.

### Bayesian phylogenetic analyses

Phylogenetic analyses of the morphological matrix were conducted in MrBayes 3.2 [41]. To model the evolution of the morphological data, we used the approach described in the seminal work by Lewis [15]. In its most basic version, his so-called ‘Mk model’ assumes that all states of a multi-state character are equally likely, as are the respective transition rates between them (we refer to this as the ‘unordered Mk’ model). An extension of this model, which was adopted from parsimony approaches, allows considering some of the characters as ‘ordered’, i.e., allows transitions only between adjacent states [42]. In a character with three states represented by labels ‘0’, ‘1’ and ‘2’, only the transitions between ‘0’ and ‘1’ and between ‘1’ and ‘2’ are allowed in an ordered character, but never directly between ‘0’ and ‘2’. As an additional extension, one can assume that there are unobserved states, i.e., states that are possible but that are not found in any of the included taxa. This might be a reasonable approach when a morphological matrix is reduced to include only part of the taxa; in such a situation, we know of possible states that a character can adopt, even though they are not realized among the taxa in our matrix. We call this the ‘full-state Mk’ model. To account for different characters evolving at different rates, we also included gamma-distributed among-character rate variation (ACRV) in the same way it is done for molecular data [among-site rate variation or ASRV, 43].

We tested the four models described above on our morphological matrix by estimating their marginal likelihoods under stepping-stone sampling [44] in MrBayes. For the ‘unordered Mk’ analysis, we made sure that the highest number used as a state label for each character corresponded to the number of states minus one (as ‘0’ is also used as a label). For the ‘ordered Mk’ analysis, we chose *a priori* those characters that can be considered to evolve in an ordered fashion; this included both characters that were originally continuous and transformed by us into discrete states (see above) and other characters that have been conceptualized in a way that suggests that they would likely adhere to a step-wise pattern of evolution (58 characters in total, see Supplementary File S3). For the ‘full-state Mk’ model, we obtained the maximum number of realized states in each character from a larger matrix at hand, which includes about twice as many ichneumonid taxa. Our data was then re-coded so that the largest state label reflects the thus estimated true size of the state space. This procedure changed the state space in 52 out of the 222 characters, in most cases only by a single state difference, but sometimes by up to three states. The R scripts to perform these matrix manipulations and the resulting morphological matrices are available as Supplementary File S4 (and as R package ‘rogue.plot‘ from CRAN). For the best-scoring model, we also ran stepping-stone sampling with equal rates among sites instead of gamma-distributed ACRV (‘no ACRV’). Stepping-stone sampling was conducted by running 50 million generations each in four independent runs under all four models, with one million generations set as an initial burn-in and then 49 steps from posterior to prior [44]. Each step consisted of one million generations, half of which were discarded as a within-step burn-in. The alpha value determining the skewness of the sampling distribution was set to 0.4 (the default value).

The preferred model as identified by the marginal likelihoods was then used for a Bayesian analysis with four independent runs, each of which with one cold and three heated chains, for 50 million generations. The ‘variable’ coding bias was invoked, as no constant characters have been scored for our dataset, but some characters showed autapomorphic states. We assessed convergence by examining plots of log likelihoods over time, ascertaining that the potential scale reduction factor (PSRF) for all scalar parameters of the model had dropped below 1.02, and recording the average standard deviation of split frequencies (ASDSF) between the four independent runs. ASDSF was below 0.015 after 50 million generations, despite the large phylogenetic uncertainty (and thus large parameter space to cover by the MCMC) introduced by the incompletely scored fossils. Half of the generations were then excluded as a conservative burn-in. All calculations were performed on UBELIX (http://www.id.unibe.ch/hpc), the HPC cluster at the University of Bern, Switzerland.

### RoguePlots to illustrate taxon placement uncertainty

To illustrate the uncertainty in the placement of fossil taxa, we developed R scripts to generate what we call ‘RoguePlots’. These plots show the posterior probabilities of the placement of a fossil on a partially resolved phylogeny, such as a consensus tree. Especially when there is a lot of phylogenetic uncertainty in a dataset, which is rather the rule than the exception in morphological data, fossils might with a non-negligible probability attach to branches that are not represented in a consensus tree (which can happen even if it is fully resolved, unless it is a strict consensus). Figure 2 illustrates how RoguePlots solve this issue: in order to represent such cases and thus ascertain that the posterior probabilities of attachments shown on a tree sum to one, the branches that are parallel to the direction of a rectangular phylogram of a RoguePlot represent fossil placements on this very branch, while the perpendicular branches stand for attachments to any unrepresented branch of which the corresponding node is the most recent common ancestor (MRCA).

**Figure 2.**
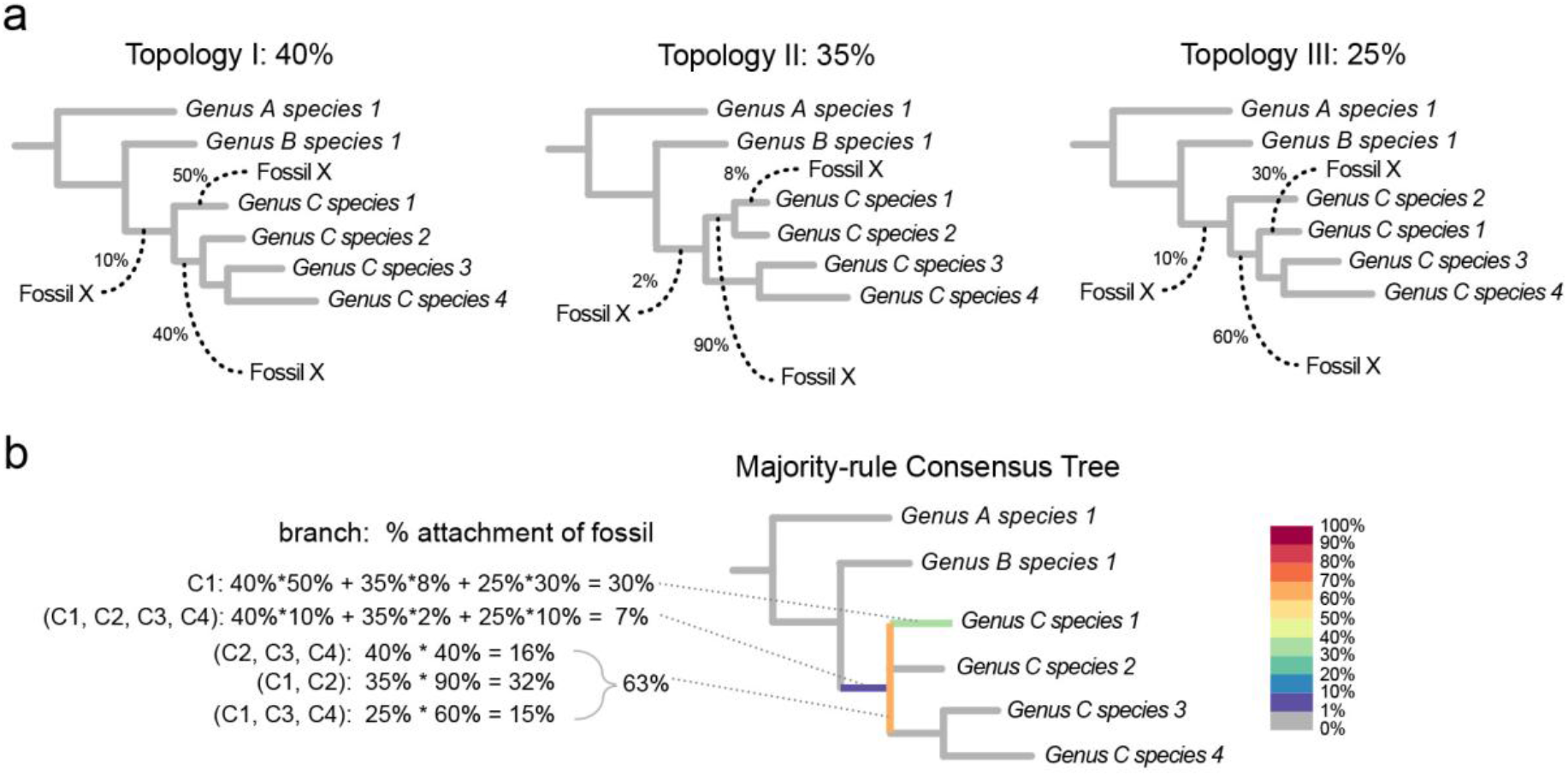
Illustration of the generation of a ‘RoguePlot’. Panel (a) shows three different topologies (I – III) as they might be found during a Bayesian MCMC analysis or based on a set of bootstrapped matrices. The frequency with which each topology is encountered is shown above. Fossil X attaches to one of three different branches in each tree. Two of these branches are also present in the majority-rule consensus tree shown in panel (b), while the third branch, respectively, is in conflict between the trees and thus appears as a polytomy in the consensus. Fossil attachment frequencies to branches present in the consensus are reflected by the colour of the branches parallel to the direction of the tree, while those to branches not present are shown on the perpendicular branch that corresponds to its most recent common ancestor. In the current example, the percentages of Fossil X to attach to a branch in the crown group of genus C sums to 93 %, while it is 7 % for it being a stem-group representative and 0 % to be with either of the other genera A and B. Classifying it as belonging to genus C is thus well-supported in this fictitious example. In addition to showing fossil placements, RoguePlots can be used to illustrate the attachment points of any taxon in a phylogeny, such as a species that was identified as a rogue.

Based on the majority-rule consensus tree from the Bayesian analysis of the morphological matrix and on 1,000 trees sampled evenly from the post-burn-in phase of the four independent runs, we generated RoguePlots to illustrate phylogenetic placements for each of the 24 fossils covered here in order to re-evaluate their genus- and higher-level classifications.

## Results

### Phylogenetic analyses

The morphological matrix has an overall coverage of 76.0 % when counting polymorphisms as informative, or 71.6 % otherwise, which corresponds to an average of 53 of the 222 characters missing per taxon. Among the extant taxa, coverage values are much higher (88.9 % and 84.5 %, respectively, depending on how polymorphisms are counted), while the fossil taxa as expected show much more missing data and thus lower coverage (20.6 % and 15.9 %, respectively). The fossil with the lowest number of characters, *Eclytus lutatus* Theobald, had 92.3 % of characters missing (only 17 of 222 characters scored), while the most completely scored fossil was *Scambus fossilobus* Spasojevic et al. [37], with ‘only‘ 67.1 % missing data (73 characters scored).

The comparison of the model likelihoods obtained from stepping-stone sampling revealed very large differences between the models, judging from Bayes factor comparisons (Table 2). While the variance in the marginal likelihoods between the four independent runs was always below six log units, the differences between the models were in the order of several hundreds. The ‘ordered Mk‘ model, which treated 58 characters as ordered (Supplementary File S3), very clearly outperformed the other models and was used for all further analyses. The ‘full-state Mk‘ model, which increased the state space of each character to that observed in other, not sampled ichneumonid taxa, reached the second-best marginal likelihood, followed by the ‘unordered Mk’ model. The model that used ordered characters but did not include among-character rate variation (‘no ACRV’), was clearly behind all other models, with a drop in likelihood of almost 850 log units (Table 2).

**Table 2.** Marginal likelihoods of the different models of morphological evolution as obtained under stepping-stone sampling.

| Model | Unordered Mk | Ordered Mk | Full-state Mk | No ACRV<sup>1</sup> |
| --- | --- | --- | --- | --- |
| Run 1 | -12,829.9 | -12,510.6 | -12,711.5 | -13,353.5 |
| Run 2 | -12,829.8 | -12,509.9 | -12,713.9 | -13,354.3 |
| Run 3 | -12,831.6 | -12,508.1 | -12,712.9 | -13,358.9 |
| Run 4 | -12,829.8 | -12,509.7 | -12,710.1 | -13,356.3 |
| Mean | -12,830.1 | -12,509.1 | -12,711.2 | -13,354.5 |
| Bayes factor<sup>2</sup> | 642.0 | - | 404.1 | 1,690.7 |

<sup>1</sup> ACRV = among-character rate variation
<sup>2</sup> Bayes factor comparison with the best model (ordered Mk). Bayes factors were calculated as $2 * (\ln L(\text{Model A}) - \ln L(\text{Model B}))$ . A Bayes factor larger than 10 is considered very strong support for Model A.

The majority-rule consensus tree of the analysis under the preferred model has a poorly resolved backbone, as can be expected from a morphology-only dataset, but recovers most of the subfamilies (Fig. 3). When rooting with *Xorides*, as supported by a previous molecular analysis that included *Xorides* as the only represenative of Xoridinae [45], the remaining subfamilies of Ichneumonidae were recovered as a monophyletic group (or in other words, the two xoridine taxa included here grouped together). We thus rooted the tree between Xoridinae and the remaining subfamilies. Higher-level relationships among subfamily groups were not resolved, despite some of these groupings having been proposed on morphological grounds (Ophioniformes, Ichneumoniformes, Pimpliformes). However, they were based mostly on larval characters, which were not included in our matrix because of extensive missing data even among extant taxa. Some of the subfamilies were not recovered either (Tryphoninae, Cylloceriinae, and Diacritinae), but most of the other subfamilies and the four tribes within Pimplinae received reasonable support. Some groups were only recovered as paraphyletic, such as Campopleginae, Delomeristini and Theronini, and the acaenitine genus *Coleocentrus* clustered with Rhyssinae. The genera *Xanthopimpla* and *Lissopimpla*, for which a separate tribe or even subfamily has been discussed previously [45], render the Theronini paraphyletic in our tree, but on a rather long branch. Overall, the tree is largely congruent with a more sparsely sampled molecular tree published recently [45], even though the latter showed better resolution along the backbone. As the goal of the current study is to re-evaluate the classification of fossil species, which relies heavily on morphology, we at this point use the morphological tree only. It allows re-assessment of most subfamily and genus placements of fossils; however, because of the low resolution of relationships between subfamilies, its value is limited when it comes to placing taxa that are backbone representatives within Ichneumonidae.

**Figure 3.**
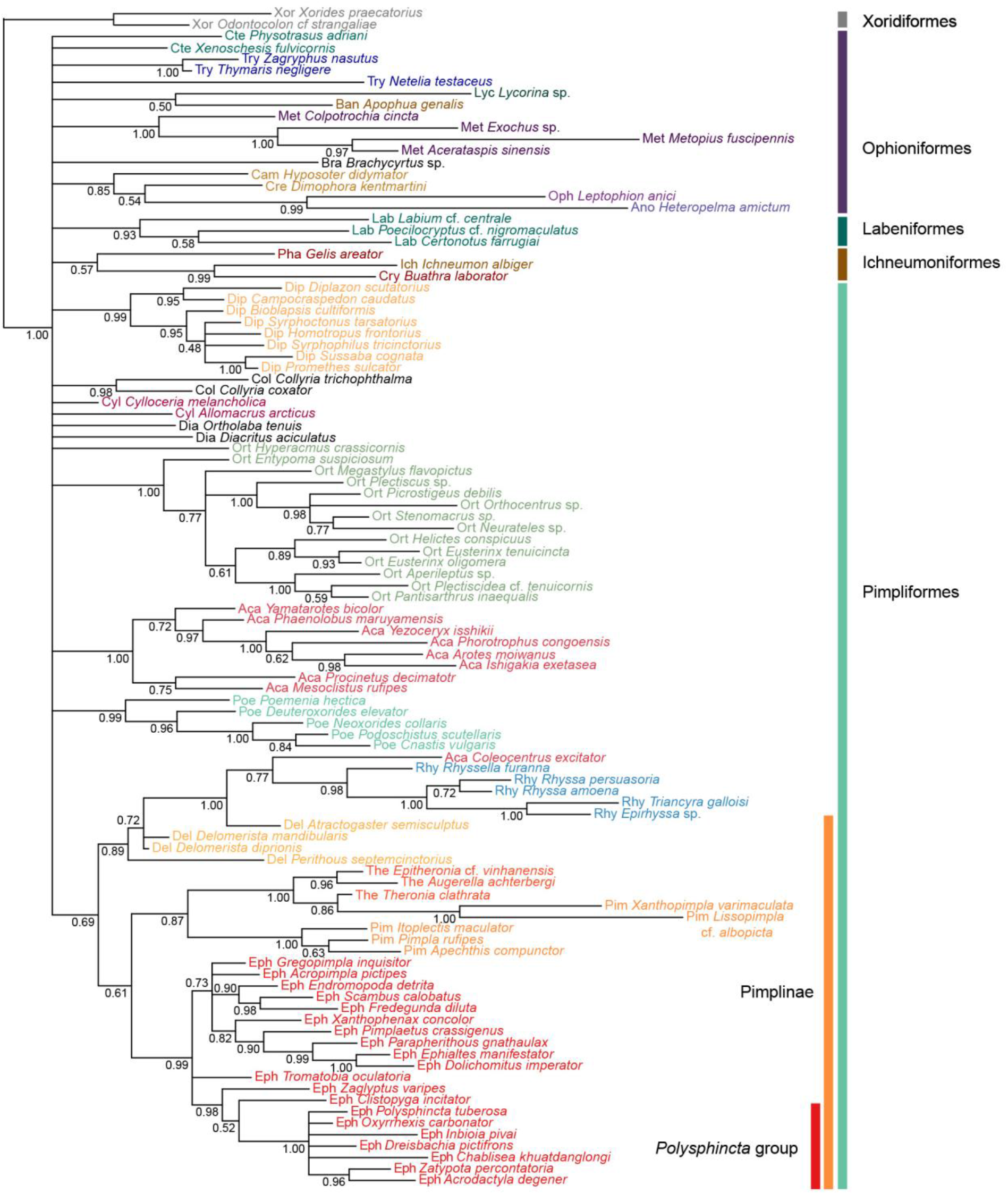
Majority-rule consensus tree of the topologies of the extant taxa sampled during the Bayesian analysis of a morphological dataset of 222 discrete characters. Values near nodes represent posterior probabilities of clades. Informal subfamily groups (Xoridiformes, Ophioniformes, etc.), the subfamily Pimplinae and the *Polysphicta* genus group (which includes parasitoids of spiders) are indicated by bars on the right. Tips are coloured according to their subfamily placement, and either subfamily or tribal (in the case of Pimplinae) classification is also indicated by the first three letters in the label as follows: Aca – Acaenitinae, Ano –Anomaloninae, Ban – Banchinae, Bra – Brachycyrtinae, Cam – Campopleginae, Col – Collyriinae, Cre – Cremastinae, Cry – Cryptinae, Cte – Ctenopelmatinae, Cyl – Cylloceriinae, Del – Delomeristini, Dia – Diacritinae, Dip – Diplazontinae, Eph – Ephialtini, Ich – Ichneumoninae, Lab – Labeninae, Lyc – Lycorininae, Met – Metopiinae, Oph – Ophioninae, Ort – Orthocentrinae, Phy – Phygadeuontinae, Pim – Pimplini, Poe – Poemeniinae, Rhy – Rhyssinae, The – Theroniini, Try – Tryphoninae, Xor - Xoridinae.

### Phylogenetic placement of 24 fossil ichneumonids

The confidence in the phylogenetic placement of the fossils (Figs 4–9) ranged from very high, with up to 99.9 % probability for attachment to a single branch (in *Xanthopimpla praeclara*, Fig. 9d), to two or three competing placements that could be either in closely related (e.g., in *Rhyssella vera*, Fig. 7d) or in unrelated groups (e.g., in *Carinibus molestus*, Fig. 4a), to very low probabilities on any individual branch (e.g., *Dolichomitus*? *saxeus*, Fig. 4d). Images of the studied fossils, along with partial RoguePlots that include all consensus tree branches that have a higher than 1 % attachment probability, are shown in Figures 4–9 (complete RoguePlots are available as Supplementary File S5). We here briefly discuss the placement of the individual fossils and, in some cases, revise the genus- and/or subfamily assignments. The fossils are sorted alphabetically according to the genus name in the original description. Taxon names are followed by the percentage of characters scored for the individual fossil species.

**Figure 4.**
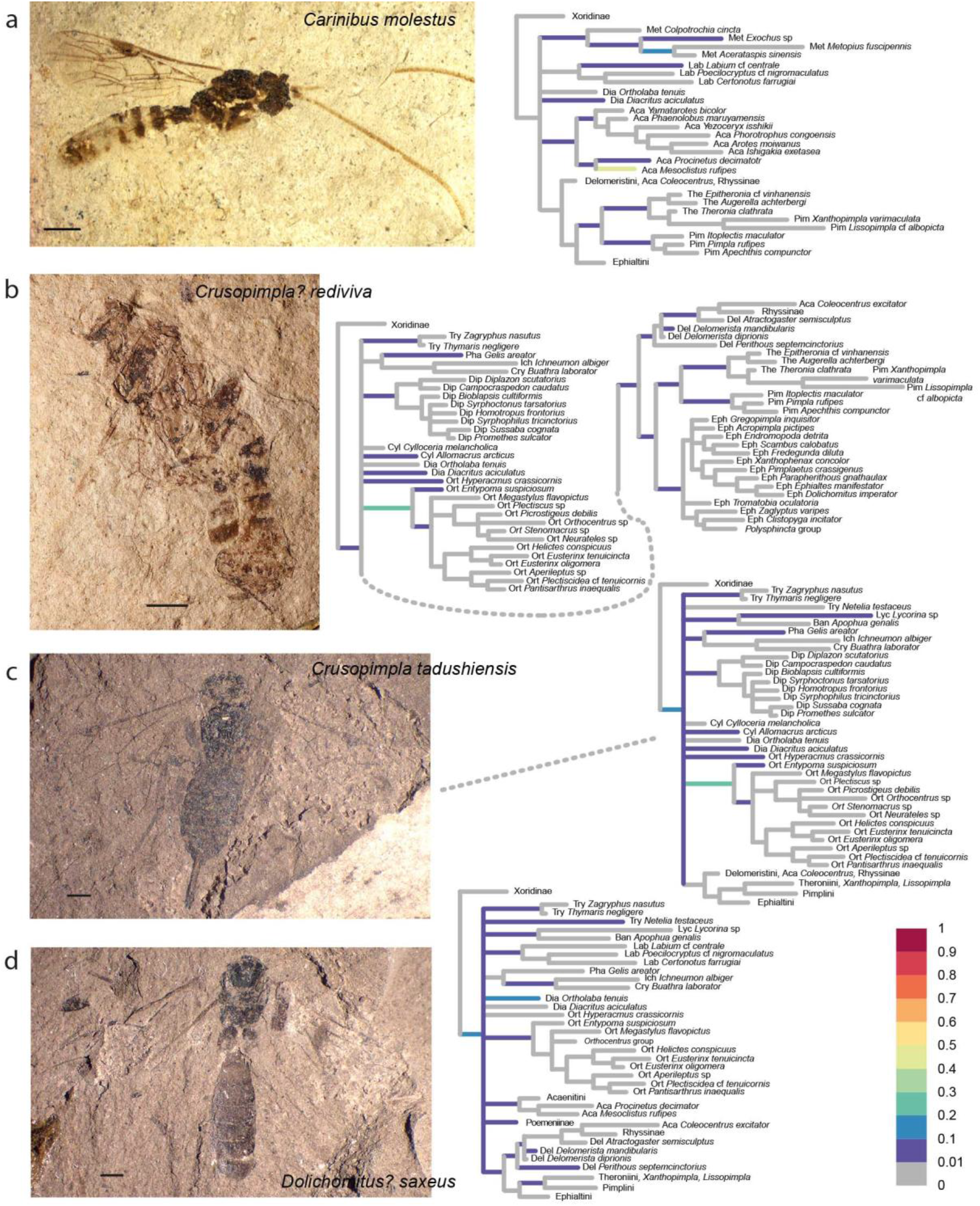
Partial RoguePlots of four fossil species including all consensus tree branches reaching more than 1 % attachment probability. Xoridinae as outgroup and in some cases subfamilies in which the fossil has been placed previously are included regardless of attachment probabilities. Branches parallel to the direction of the phylogram are coloured according to the probability of direct attachment of the fossil to the branch, while perpendicular branches represent the sum on any branch not in the consensus tree, of which the corresponding node is the most recent common ancestor. The scale bar in the photographs represents 1 mm. a) *Carinibus molestus*, holotype USNM 580881. b) *Crusopimpla rediviva*, holotype #2156, ©President and Fellows of Harvard College. c) *Crusopimpla tadushiensis*, holotype #PIN 3364/277, ©Russian Academy of Sciences, Moscow. c) *Dolichomitus*? *saxeus*, holotype #PIN 3364/31, ©Russian Academy of Sciences, Moscow.

**Figure 5.**
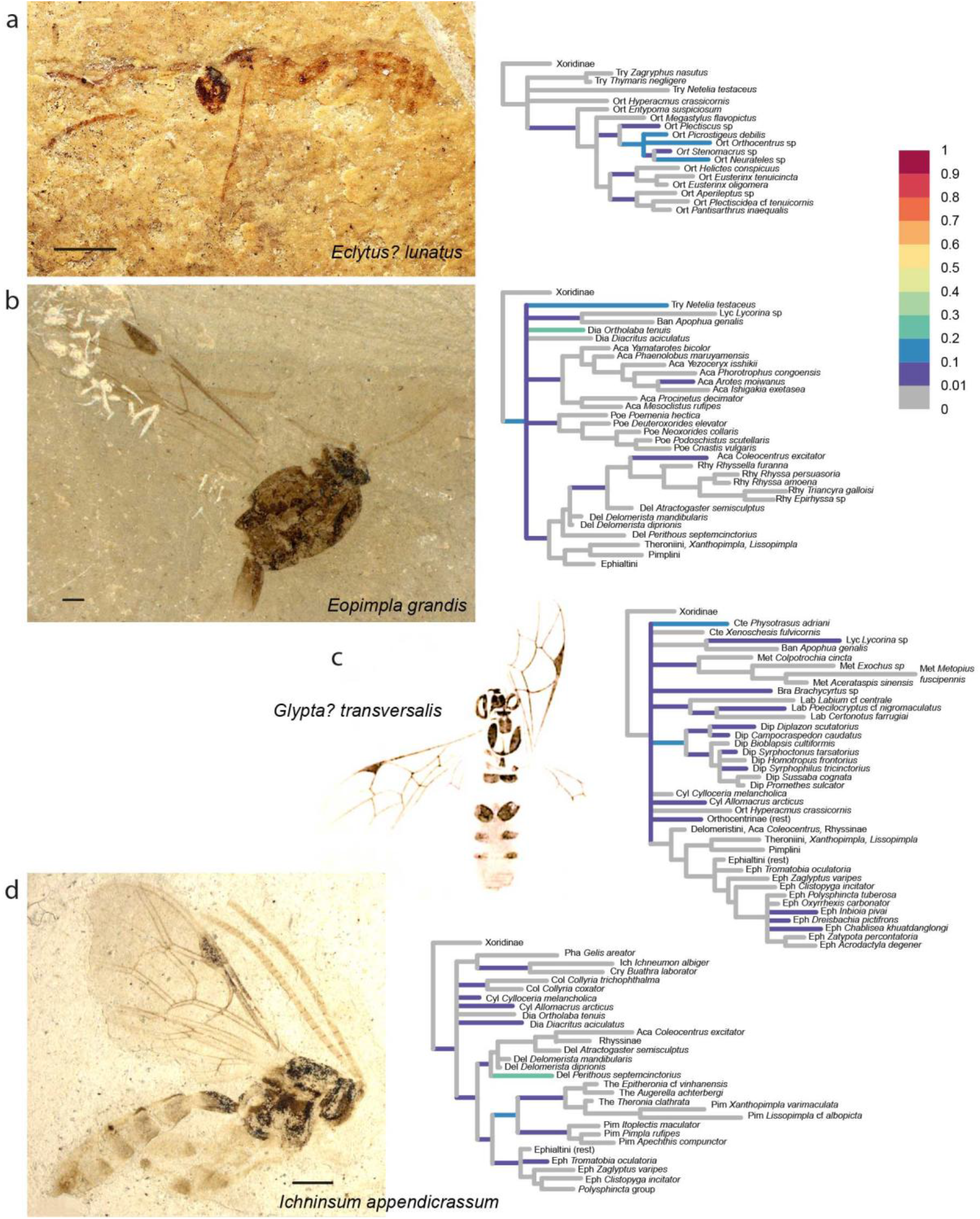
Partial RoguePlots of four fossil species including all consensus tree branches reaching more than 1 % attachment probability. Xoridinae as outgroup and in some cases subfamilies in which the fossil has been placed previously are included regardless of attachment probabilities. Branches parallel to the direction of the phylogram are coloured according to the probability of direct attachment of the fossil to the branch, while perpendicular branches represent the sum on any branch not in the consensus tree, of which the corresponding node is the most recent common ancestor. The scale bar in the photographs represents 1 mm. a) *Eclytus*? *lutatus*, holotype PALE-1418, © President and Fellows of Harvard College. b) *Eopimpla grandis*, holotype USNM 66581, © Smithsonian Institute. c) *Glypta transversalis*, modified from original drawing [Plate X, Fig. 25 in 47]. d) *Ichninsum appendicrassum*, holotype UCM 39378.

**Figure 6.**
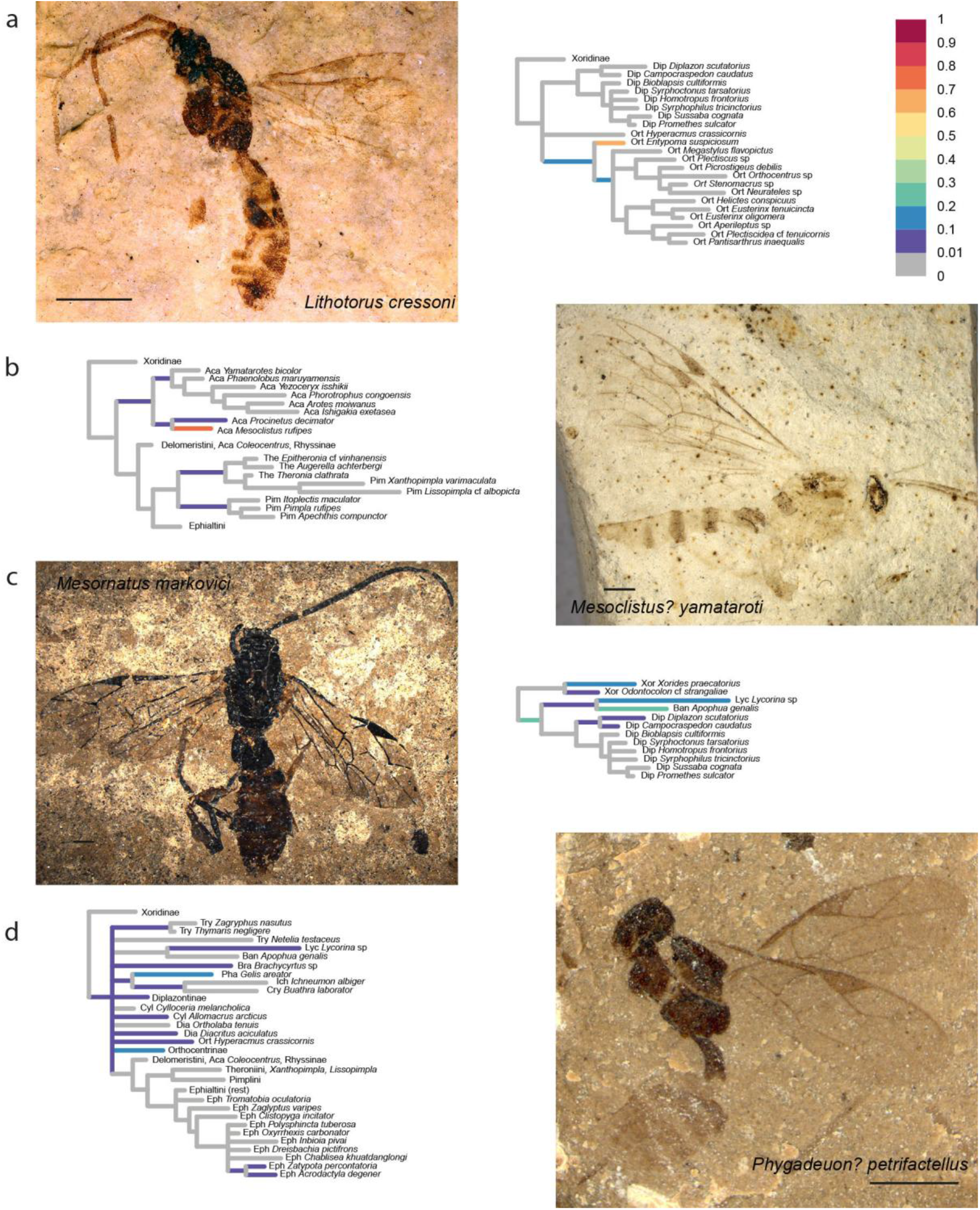
Partial RoguePlots of four fossil species including all consensus tree branches reaching more than 1 % attachment probability. Xoridinae as outgroup is included regardless of attachment probabilities. Branches parallel to the direction of the phylogram are coloured according to the probability of direct attachment to the branch, while perpendicular branches represent the sum on any branch not in the consensus tree, of which the corresponding node is the most recent common ancestor. The scale bar in the photographs represents 1 mm. a) *Lithotorus cressoni*, holotype PALE-4652, © President and Fellows of Harvard College. b)*Mesoclistus*? *yamataroti*, holotype UCM 62725. c) *Mesornatus markovici*, holotype SF MeI 15245. d) *Phygadeuon*? *petrifactellus*, holotypeUSNM 66580 © Smithsonian Institute.

**Figure 7.**
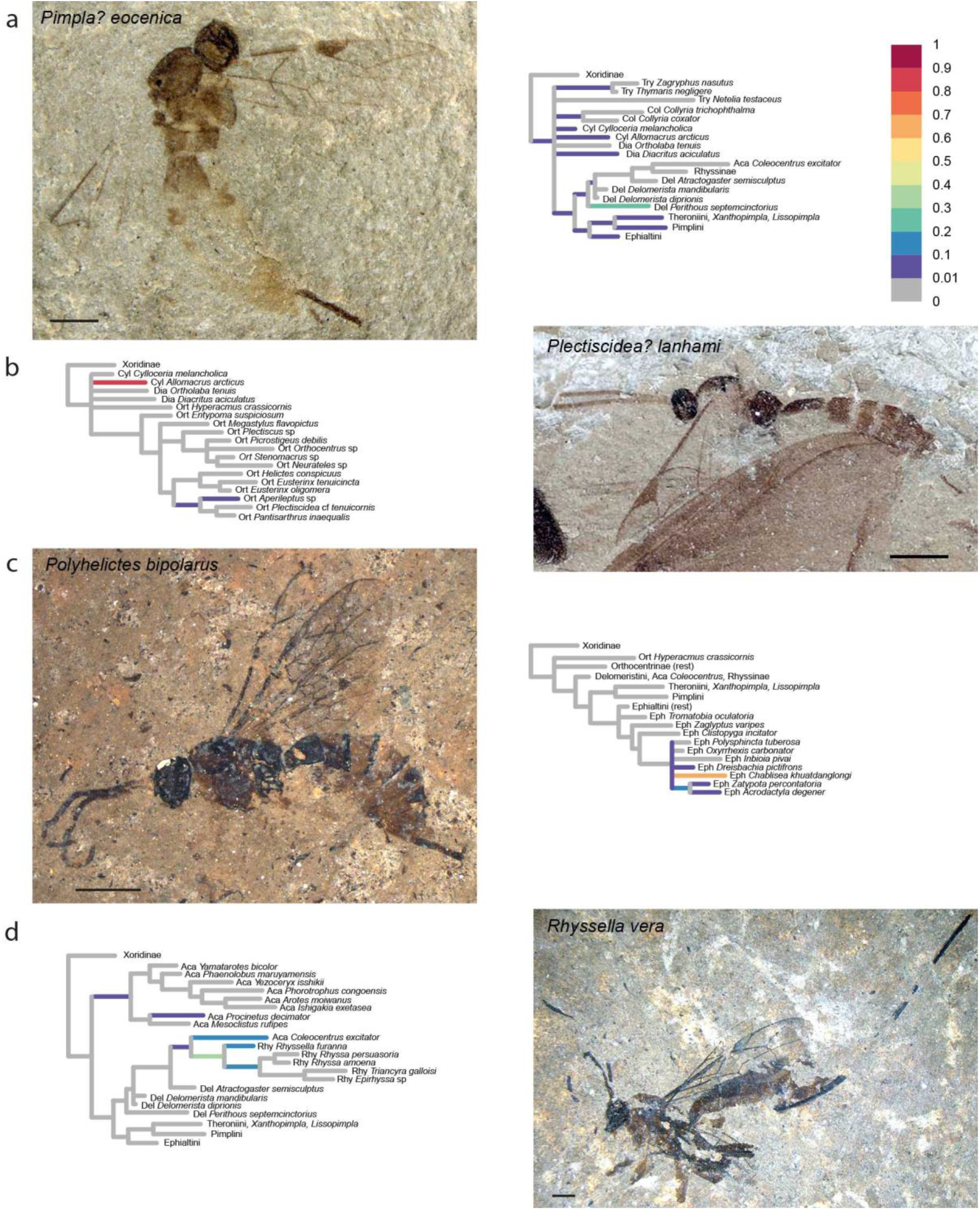
Partial RoguePlots of four fossil species including all consensus tree branches reaching more than 1 % attachment probability. Xoridinae as outgroup is included regardless of attachment probabilities. Branches parallel to the direction of the phylogram are coloured according to the probability of direct attachment to the branch, while perpendicular branches represent the sum on any branch not in the consensus tree, of which the corresponding node is the most recent common ancestor. The scale bar in the photographs represents 1 mm. a) *Pimpla*? *eocenica*, holotype USNM 66582, © Smithsonian Institute. b) *Plectiscidea*?*lanhami*, holotype UCM 19167, © University of Colorado Museum of Natural History. c) *Polyhelictes bipolarus*, holotype MeI 16069. d) *Rhyssella vera*, holotype MeI 8814.

**Figure 8.**
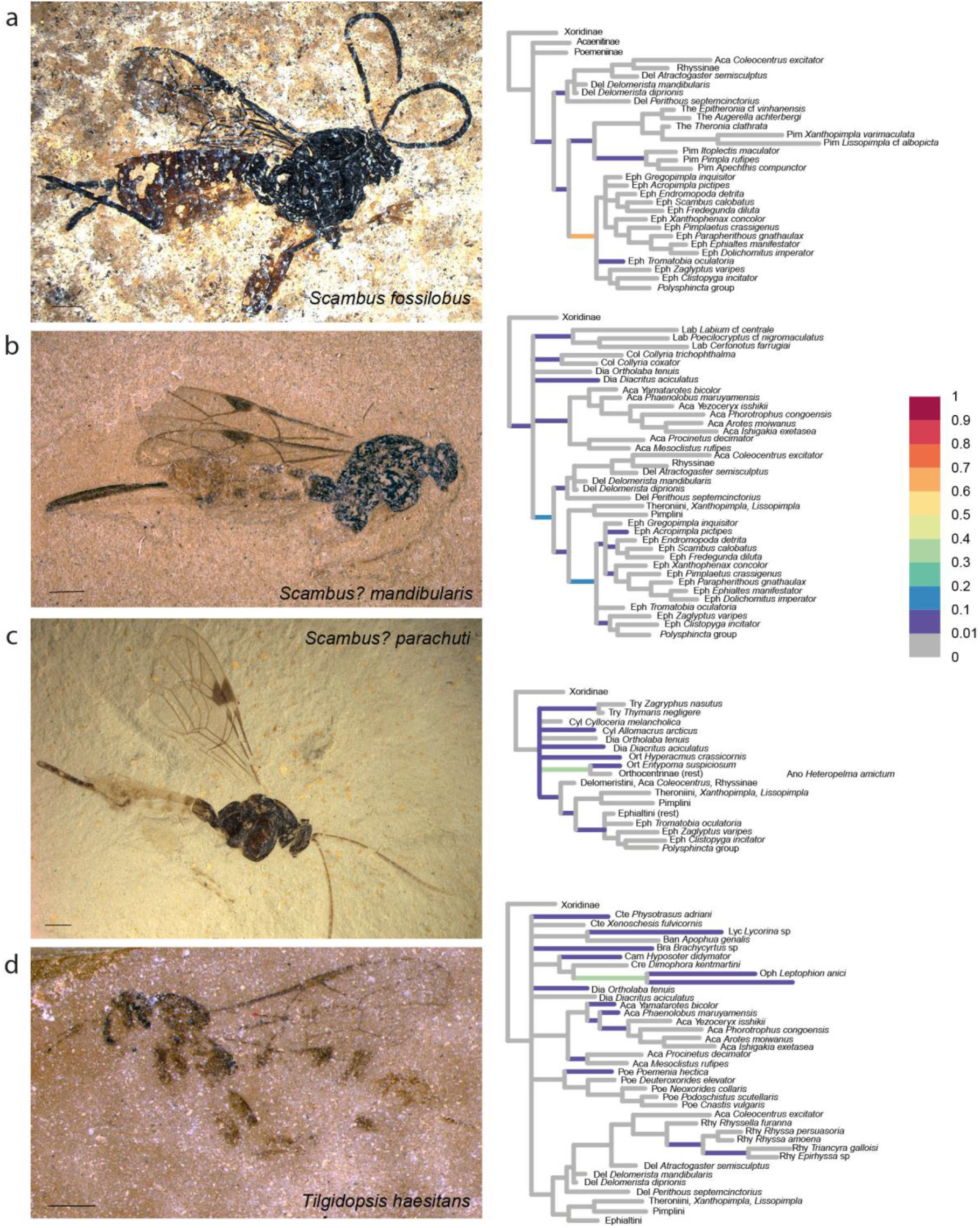
Partial RoguePlots of four fossil species including all consensus tree branches reaching more than 1 % attachment probability. Xoridinae as outgroup is included regardless of attachment probabilities. Branches parallel to the direction of the phylogram are coloured according to the probability of direct attachment to the branch, while perpendicular branches represent the sum on any branch not in the consensus tree, of which the corresponding node is the most recent common ancestor. The scale bar in the photographs represents 1 mm. a) *Scambus fossilobus*, holotype SF MeI 13431. b) *Scambus*? *mandibularis*, holotype USNM 501474. c) *Scambus*? *parachuti*, holotype 565885. d) *Tilgidopsis haesitans*, holotype USNM 66931, © Smithsonian Institute.

**Figure 9.**
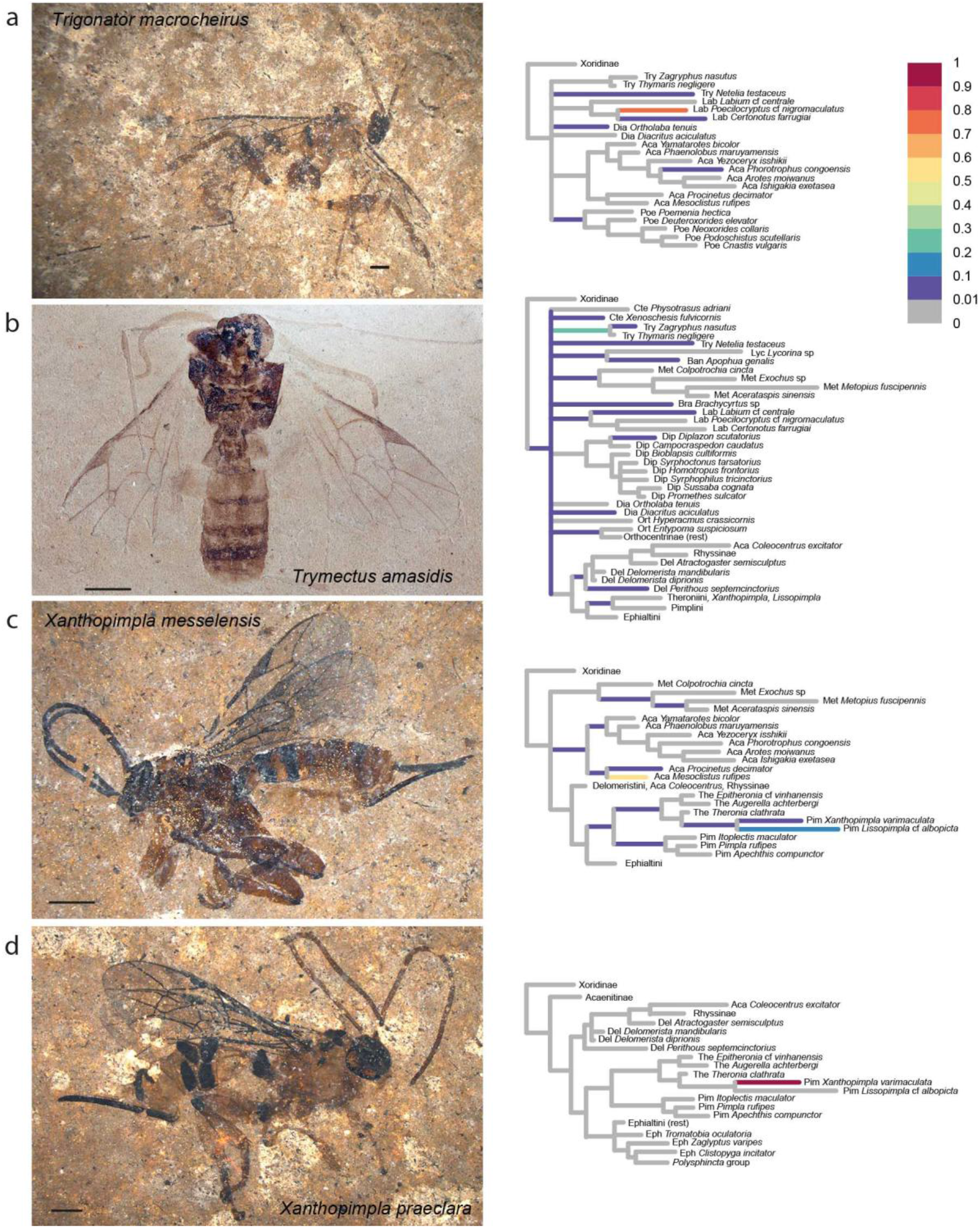
Partial RoguePlots of four fossil species including all consensus tree branches reaching more than 1 % attachment probability. Xoridinae as outgroup is included regardless of attachment probabilities. Branches parallel to the direction of the phylogram are coloured according to the probability of direct attachment to the branch, while perpendicular branches represent the sum on any branch not in the consensus tree, of which the corresponding node is the most recent common ancestor. The scale bar in the photographs represents 1 mm. a) *Trigonator macrocheirus*, holotype SF MeI 17304. b) *Tryphon amasidis*, holotype UCM 15690, © University of Colorado Museum of Natural History. c) *Xanthopimpla messelensis*, holotype SF MeI 16988. d) *Xanthopimpla praeclara*, holotype SF MeI 17300.

*Carinibus molestus* Spasojevic et al. [7] – 22 %, Figure 4a. This taxon was described in its own genus with uncertain subfamily placement. We can confirm this notion but can narrow down potential subfamilies: The most likely placement in our analysis is with the acaenitine *Mesoclistus* (42%), and the total probability for it to be an acaenitine sums to 55%. But there is an alternative placement in the subfamily Metopiinae (total 32%), with a probability so high that it cannot be ignored. Given the unique combination of characters described by the authors [7], the genus might well represent an ancestor or extinct subfamily; we thus leave the genus in *incertae subfamiliae*.

*Crusopimpla*? *rediviva* [27,28] – 18 %, Figure 4b. This species attaches to various places in the tree, as does the type species of the genus, *C. tadushiensis* (see next fossil), with the exception that a placement as one of the more ancestral lineages within Pimplinae received some support (22 %). The placement on the stem branch of Orthocentrinae was deemed more probable, however (25 %), but due to the possibility of an artefact due to model misspecification (see discussion section), we leave it in its current position.

*Crusopimpla tadushiensis* Kopylov et al. [28] – 17 %, Figure 4c. This species appears all over the tree in our analysis, with placements in seven different subfamilies and on the ancestral branch of Ichneumonidae minus Xoridinae, but never in the subfamily Pimplinae in which it was originally described. This genus was erected as a stem-lineage representative of the subfamily. Given the possibility that model misspecification caused this difference in interpretation, and that the preferred placement in Orthocentrinae (22 %) seems highly unlikely given the invariably smaller size of its representatives, we refrain from suggesting any changes to its classification.

*Dolichomitus*? *saxeus* Kopylov et al. [28] – 14 %, Figure 4d. The placement of this taxon was highly uncertain, which might be expected given its poor preservation, but there seems a clear tendency for it to take up a basal position. This might reflect a true ancestral placement of the fossil or could be due to ‘stem-ward slippage’ [46], the phenomenon that incompletely preserved fossils tend to attach closer to the root than would be accurate (see also discussion section). The sparse character evidence precludes any clear assessment of this question, and as a placement within Pimplinae could not be ruled out either (17 %), and genus assignment was already marked with a question mark by the authors of the original description, we leave it in its current taxonomic position.

*Eclytus*? *lutatus* Scudder [47] – 8 %, Figure 5a. Even though it was described in a tryphonine genus, this fossil received very high support in our analysis to belong to Orthocentrinae (95.2 %), and there-in to the *Orthocentrus*-group of genera (85.3 %). All placements on other branches in the tree, including anywhere near the three Tryphoninae taxa we included, had less than 1 % frequency. The fossil is not very well preserved and its placement difficult, as pointed out previously [7,48]. Furthermore, we did not manage to include an extant member of the genus *Eclytus* in our analysis, and this high placement probability might be due to insufficient taxon sampling and/or size-related homoplasy of the scored morphological characters (see discussion). We thus agree with the decision taken by Spasojevic et al. [7] and leave it as questionable within its original genus. Future analyses might include more taxa from the morphologically quite diverse Tryphoninae and thus allow for a more meaningful conclusion. However, a placement in Orthocentrinae should definitely be considered, given the similarities to some genera of that subfamily in terms of the small size and shape of the first tergite.

*Eopimpla grandis* Cockerell [49] – 16 %, Figure 5b. As stated previously [7], there is little evidence for placing this fossil in Pimplinae, and the phylogenetic analysis was highly ambiguous about subfamily placement. The fossil clustered with 22.5 % with *Ortholaba* (Diacritinae), 19 % with *Netelia* (Tryphoninae), and 16.5 % on the ancestral branch leading to Ichneumonidae without Xoridinae. Removing it from Pimplinae into an uncertain subfamily association [7] thus seems justified.

*Glypta*? *transversalis* Scudder [47] – 12 %, Figure 5c. Spasojevic et al. [7] suggested that this fossil does not fit very well in Banchinae, but instead could be associated with Lycorininae or with the genus *Physotarsus* in Tryphoninae. While the fossil was never recovered with the banchine in our dataset (*Apophua*, a genus closely related to *Glypta*), it indeed clustered most often with *Physotarsus* (13%), but there were also numerous alternative placements that could not be ruled out. In lack of a well-supported alternative, we leave the fossil in its original placement, but follow Spasojevic et al. [7] in adding a question mark behind the genus name.

*Ichninsum appendicrassum* Spasojevic et al. [7] – 26 %, Figure 5d. The authors of this genus avoided subfamily placement because of presumably plesiomorphic character combinations. Our results here confirm what they suggested as the most likely subfamily placement (Pimplinae) with quite high cumulative posterior probability (65.3 %). Even though the fossil cannot be associated with a single tribe within the subfamily, we move it from *incertae subfamiliae* to Pimplinae.

*Lithotorus cressoni* Scudder [47] – 20 %, Figure 6a. This fossil was described in the subfamily Diplazontinae, but both Townes [50] and Spasojevic et al. [7] mentioned the possibility that it is instead closely related to the *Helictes*-group of genera in Orthocentrinae. Our results confirm this notion and place it either as a sister taxon to *Entypoma* or as a stem-lineage representative of the subfamily (without *Hyperacmus*). The sum of placement probabilities in Orthocentrinae approaches 98.5 %, while the original placement in Diplazontinae is not supported at all. We thus move the genus *Lithotorus* to Orthocentrinae.

*Mesoclistus*? *yamataroti* Spasojevic et al. [7] – 23 %, Figure 6b. We can confirm the subfamily placement here (Acaenitinae: 96%), and indeed observed the highest probability of it clustering with *Mesoclistus* (77%). As we have only included two other genera of the *Coleocentrus* genus group in this analysis (*Coleocentrus* and *Procinetus*), we leave the genus placement as uncertain, as it was already suggested in the original description.

*Mesornatus markovici* Spasojevic et al. [37] – 27 %, Figure 6c. This taxon is placed with 23 % probability on the stem lineage leading to all ichneumonids except Xoridinae, and with 20 %, 15 %, and 14.5 %, respectively, with *Apophua* (Banchinae), *Lycorina* (Lycorinae), and *Xorides* (Xoridinae). We can thus confirm the uncertain subfamily placement of this genus; it might even belong to a stem lineage or now extinct subfamily.

*Phygadeuon*? *petrifactellus* Cockerell [49] – 14 %, Figure 6d. Spasojevic et al. [7] added a question mark to the genus assignment, but left the taxon within the then subfamily Cryptinae. As this subfamily has in the meantime been split by Santos [51], the correct placement would now be in the subfamily Phygadeuontinae. Our results largely agree with this placement, but emphasize even more the large uncertainty: it grouped with *Gelis areator* 18 % of the time, but was also found as a stem-lineage representative of Orthocentrinae in 14 % of the trees. We leave the current classification unchanged but emphasize the uncertainty of the placement.

*Pimpla*? *eocenica* Cockerell [52] – 16 %, Figure 7a. Spasojevic et al. [7] revised this fossil and added a question mark to the genus assignment, citing high uncertainty in this genus placement due to poor preservation of the fossil. Our morphological analysis confirmed the high uncertainty in the placement, which attached on several mostly rather basal branches within Pimplinae, but also with other subfamilies, most of all Diacritinae, Collyriinae, and Cylloceriinae. Summing the probabilities of it associating with pimpline taxa revealed a slight preference of this over any other subfamily (56 %); within the subfamily, *P. eocenica* was most often ending up in the tribes Delomeristini, Pimplini and Theroniini or on the branches close to the ancestor of the subfamily. The highest probability (23 %) was for *P. eocenica* to be sister to *Perithous septemcinctorius*, a placement not supported by any particular character. We thus leave *P. eocenica* in Pimplinae and, with a question mark, in the genus *Pimpla*.

*Plectiscidea lanhami* Cockerell [53] – 13 %, Figure 7b. Spasojevic et al. [7] noted uncertainty in the generic placement of this fossil, mostly due to the pentagonal areolet which does not occur in any of the recent representatives of the genus. But they agreed with the placement in Orthocentrinae. Our analysis now recovers the fossil with 89 % with the very small genus *Allomacrus*, and only with about 7.5 % in Orthocenrinae. However, *Allomacrus* was variously classified within either Cylloceriinae or Orthocentrinae and these subfamilies are certainly closely related. Even though *Allomacrus* has an open areolet, its shape indicates that it might be pentagonal when closed. As far as visible in the fossil, the wing venation (i.e., proportions of pterostigma and fore wing cells), humped first tergite and proportions of the remaining tergites, and possibly long ovipositor are indeed very similar to *Allomacrus*, thus we transfer the fossil to this genus: *Allomacrus lanhami*, comb. nov.

*Polyhelictes bipolarus* Spasojevic et al. [37] – 26 %, Figure 7c. The authors of this genus and species mentioned that the character combination would allow placement in two not very closely related groups: the *Helictes* genus group in Orthocentrinae, or the spider parasitoids of the *Polysphincta* genus group in the pimpline tribe Ephialtini. This ambiguity is expressed both in the genus and species names of the taxon. However, our analysis places it very firmly in the *Polysphincta* group (100 %). This somewhat surprising result is probably due to some measurement characters in the fore wing that were not taken into account when the taxon was described, such as the rather elongate radial cell and relatively long vein r-rs in the forewing, which are consistently short in Orthocentrinae.

*Rhyssella vera* Spasojevic et al. [37] – 27 %, Figure 7d. This fossil received highest probability for attaching to the stem lineage of the Rhyssinae taxa we sampled (36 %), followed by being sister to the extant *Rhyssella furanna* (18 %) or on the branch leading to the remaining Rhyssinae (17 %). There was some probability also that it clustered with Acaenitinae (total 24 %), mostly with *Coleocentrus* (12 %), which was recovered here as the sister group to Rhyssinae. Nevertheless, we consider the original placement as being supported by this analysis (total in Rhyssinae: 72 %), with the genus *Rhyssella* as the best option.

*Scambus fossilobus* Spasojevic et al. [37] – 33 %, Figure 8a. Our analysis recovers this taxon with high certainty within Pimplinae (91 %), where the highest probability attaches it to the stem- of the tribe Ephialtini (67%). This confirms the author’s claim of the first unequivocal representative of the subfamily, and the genus placement, even though not directly confirmed here, cannot be refuted, as we did not adequately sample the diversity of the large genus *Scambus*.

*Scambus*? *mandibularis* Spasojevic et al. [7] – 20 %, Figure 8b. This fossil also was placed mostly on stem lineages in various subfamilies, but with most of the weight in Pimplinae (sums to 53%), most likely either as a stem lineage representative or ancestral to the tribe Ephialtini. Genus placement is not resolved though, and we thus leave it as uncertain within *Scambus*.

*Scambus*? *parachuti* Spasojevic et al. [7] – 29 %, Figure 8c. This taxon received a placement in various subfamilies, but mostly in a very basal position or even as a stem lineage. This is also the case for the subfamily Pimplinae in which it was described. The highest probability is observed for a stem-lineage placement in Orthocentrinae, but this is not decisive (36 %). Even though the placement in Pimplinae is not supported here, it cannot be rejected either (sums to 8 %). We thus leave it as uncertain in the genus *Scambus*.

*Tilgidopsis haesitans* Cockerell [54] – 11 %, Figure 8d. Described in Ophioninae but with reported similarities to the unrelated Poemeniinae, Cockerell already made it clear that subfamily placement is not simple in this taxon. Spasojevic et al. [7] formalized this by moving it to *incertae subfamiliae*. Interestingly, our analyses put it closer to Ophioninae again, but most likely as a stem lineage with 37.5 % probability to attach to the branch leading to *Leptophion* (Ophioninae) and *Heteropelma* (Anomaloninae). Other placements in various subfamilies (Acaenitinae, Brachycyrtinae, Diacritinae, Ctenopelmatinae, etc.) cannot be fully excluded either.

*Trigonator macrocheirus* Spasojevic et al. [37] – 23 %, Figure 9a. Despite rather sparse sampling of the subfamily Labeninae, this fossil is firmly placed within that subfamily (87 %), with closest ties to *Poecilocryptus* (80 %) and *Certonotus* (6 %), the sole representatives of the two tribes that the authors associated it with. However, without a denser sampling of the labenine genera, tribal placement of the genus cannot be made with any certainty.

*Trymectus amasidis* [55] – 22 %, Figure 9b. This fossil was originally described in *Tryphon* and is rather well preserved; the more surprising was its unresolved placement. It ended up on the branch leading to the Tryphoninae taxa *Zagrphus* and *Thymaris* in 24% of the trees, with the rest of the probabilities distributed rather evenly among numerous taxa in subfamilies as different as Metopiinae, Ctenopelmatinae, Banchinae, Pimplinae etc., or on some of the most basal branches in the tree. The placement as a new genus *Trymectus* with uncertain subfamily placement by Spasojevic et al. [7; named after the first letters of the first three afore-mentioned subfamilies] can thus here be confirmed.

*Xanthopimpla messelensis* Spasojevic et al. [37] – 30 %, Figure 9c. While the similarities of this fossil with recent representatives of the genus seem striking at first glance [see Fig. 3 in 37], our analysis placed it with higher probability in Acaenitinae (50 % with *Mesoclistus*), even though *Xanthopimpla* and its sister genus *Lissopimpla* remain a clear possibility (19 % in total). We acknowledge that we judged some of the characters typical of *Xanthopimpla* as not clear enough for scoring in the fossil, even though they are indicated, e.g., the groves on the metasomal tergites and enlarged claws. Adding these characters and achieving a better coverage of the morphological diversity among extant *Xanthopimpla* species would probably tip the scale in favour of the original placement. The shape and length of the ovipositor and absence of a large, triangular hypopygium strongly contradict a placement in Acaenitinae. We thus refrain from making any formal subfamily changes here, but put a question mark behind the genus name to express the uncertainty in the phylogenetic placement. Especially, the fossil could also belong to a stem lineage of *Xanthopimpla* + *Lissopimpla* or of Theronini.

*Xanthopimpla praeclara* Spasojevic et al. [37] – 29 %, Figure 9d. In contrast to the previous fossil, this species was placed very firmly (99.9 %) with the recent *Xanthopimpla varimaculata*, confirming that this genus dates back to the Early to Mid Eocene.

## Discussion

### Evolutionary models for morphological data

The application of model-based approaches to a specific type of character requires the availability of sufficiently realistic models of its evolution, and the discussion whether the simplifying Mk model [15] is suitable to infer phylogenetic relationships from discrete, morphological data is still on-going. It was further sparked by a simulation study suggesting the superiority of the Bayesian approach over parsimony for morphological data [56], but the specifics of the simulation procedure, especially the single set of relative branch lengths used, are still under discussion [57,58].

Even though a lot remains to be done in the field of morphology models, the stochastic framework allows for a direct comparison of the fit of different models, as we have conducted here when comparing the unordered versus ordered Mk models and the expansion of the state space as implied by a larger morphological matrix (Table 2). The very large differences in model likelihoods revealed by stepping-stone sampling suggest that provided with a large-enough data matrix (222 characters times 127 taxa), such comparisons are very instructive. In our case, the largest effect was observed when comparing the ‘ordered Mk’ model with or without gamma-distributed among character rate variation (ACRV), an observation made earlier for molecular data [59]. ACRV will likely constitute a vital part of any morphology model, as was already suggested by Lewis [15], but systematic reviews of empirical matrices are currently missing. The second-ranked Bayes factor resulted from the comparison of ‘unordered’ versus ‘ordered’ data, with characters for ordering chosen *a priori* according to character conceptualization (Supplementary File S3). Some of the ‘ordered’ characters (13 out of 58) in fact represent discretised continuous characters, and it remains to be shown whether character ordering is vital also in datasets where continuous characters are analysed as such [60]. Finally, we tested whether increasing the state space of each character by adding unobserved states as they appeared in a larger matrix of ichneumonid taxa (unpublished data). Even though one might argue that such an approach leads to a more realistic morphology model, it received lower marginal likelihoods in our analyses (Table 2). A possible explanation of this is that we used a fully symmetrical model, where transitions between different states and thus also state frequencies are assumed to be equal at stationarity. Adding states that are not actually observed among the data creates additional asymmetry in state frequencies, which might be the reason for the lower marginal likelihood of that model.

We also attempted to run an asymmetric model which allows state frequencies to be unequal following a beta distribution for two-state or Dirichlet distribution for multi-state characters. This extension to the model has already been suggested by Lewis [15] and is implemented in MrBayes [41]. It has been demonstrated recently to outperform the symmetric model in about half of 206 empirical datasets, most of which were rather small [14]. On our data matrix of 222 characters times 127 taxa, even using a fixed level of asymmetry instead of a hyper-prior, we found computation times to be prohibitive (more than 1,100 CPU hours for 10 million generations).

### Bayesian phylogenetic fossil placement

Our morphological phylogenetic analysis exemplifies the use of Bayesian inference in the classification of fossil taxa and shows how placement uncertainty can be illustrated using RoguePlots. The advantages of reproducible, stochastic analyses of fossil placement have been discussed previously [16,61], and we here further emphasize the benefits of the direct assessment of alternative placements that comes with Bayesian approaches. Placing fossils in a classification system that is strongly biased towards extant taxa, as it is the case in most groups with a poor (or at least poorly studied) fossil record, triggers difficulties that go beyond incomplete fossil preservation. Representatives of a stem lineage might not yet possess all the synapomorphies of the crown group, requiring an extension of the circumscription of a taxon. And even more important, because the divergence times for most taxa remain poorly known or contentious, it is often unclear to what degree a fossil species can be placed at all in a certain taxonomic rank: the genus, tribe, or even family in question might not have existed yet at the time the species fossilized. A phylogenetic perspective on fossil classification is thus vital for a proper evaluation and communication of both the position of a fossil in the taxonomic system and the amount of uncertainty associated with it.

RoguePlots provide a natural way of representing placement uncertainty on a tree summary like a consensus tree, but also on individual trees, such as a maximum likelihood tree or shortest tree under the parsimony criterion. The approach can take sets of trees from a Bayesian analysis, but also those resulting from bootstrapped data matrices; it illustrates the frequencies with which different fossil placements occur among the different trees. One difficulty arising from the need to represent relationships among fossil and extant taxa in a potentially very heterogeneous set of topologies lies in the restrictions imposed by using a single tree plot: even if the tree in question is fully resolved, some of the trees in the set will most likely contain clades not represented in it. RoguePlots circumvent this issue by making use of the distinction in branches that are parallel versus perpendicular to the direction of a rectangular phylogram: parallel branches are used to illustrate the probability of direct attachment to the branch, while perpendicular branches represent the sum on any branch not in the tree, of which the corresponding node is the most recent common ancestor. This approach ascertains that the shown placement probabilities sum to one, which might not be the case otherwise [e.g., see Fig. 8 in 23]. While it is possible to produce RoguePlots based on trees obtained from a maximum likelihood or parsimony analysis of a morphological dataset, their interpretation is much more intuitive in the Bayesian context, where the frequency of a clade in the set of trees represents its posterior probability. In contrast, bootstrapping a morphological matrix will necessarily remove some characters, while multiplying others, precluding a straight-forward interpretation of the resulting placement frequency.

### Morphological phylogenetics versus character evidence for fossil placement

Using a stochastic framework for the phylogenetic analysis of fossil taxa does neither replace nor preclude careful character evaluation, but instead is based on it being made explicit. The step of coding morphological observations into discrete characters and states is crucial for this endeavour and certainly comes with a string of pitfalls, from character conceptualization to the delimitation of character states. In the ideal case, characters that would have been used as arguments to place a fossil can be adequately represented in a morphological matrix, in which case the phylogenetic reconstruction method identifies them as characters that evolve at a low rate (which is nearly equivalent to identifying them as synapomorphies). They will lead to a high probability of the predicted placement of the fossil, or at least to an accurate representation of the uncertainty of this placement (but see below). In our analyses, this was the case for the vast majority of fossils, where the RoguePlots provided matching illustrations for alternative placements that had already been discussed using character evidence in the original descriptions or revisions of the fossils [7,28,37]. In several cases, the morphological phylogenetic analysis helped identify informative characters that have not been used much previously to define higher taxa, and were thus omitted from consideration in the original descriptions. The best example here is *Polyhelictes bipolarus*, a taxon originally described with uncertain subfamily association [37], as the traditionally used characters would imply a placement either in the *Helictes*-group of genera in the subfamily Orthocentrinae, or among the spider parasitoids of the *Polysphincta*-group in Pimplinae. Examination of the morphological matrix showed that the unequivocal placement in the latter group in the morphological phylogeny was driven mostly by measurements of forewing veins and cells, characters not typically used in ichneumonid subfamily diagnoses. A phylogenetic analysis can also expose overconfidence in fossil placements that are based on individual characters by identifying alternative placements. This was probably the case with *Rhyssella vera*, which was initially described in the recent genus based on the strongly petiolate areolet (Spasojevic et al. 2018), while our analysis suggests a more basal placement as a stem lineage rhyssine.

On the other hand, a phylogenetic analysis is only as good as the underlying morphological matrix, and generating the latter does not come without its pitfalls. We below discuss five issues that can result in erroneous phylogenetic placement of fossils: limitations in character sampling, limitations in taxon sampling, artefacts caused by the fossilization process, limited phylogenetic signal in the morphological data, and misspecification of the evolutionary model. The identification of resulting artefacts is crucial when interpreting the results of a morphological phylogenetic analysis and can point to way to improving the representation of the morphological diversity of extant and fossil taxa in the matrix in the future.

#### Insufficient character sampling

Probably the most common cause of differences between the *a priori* assessment of the placement of a fossil and that resulting from a phylogenetic analysis are differences in the representation of character evidence. While the coding of some discrete characters is rather straight-forward, such as the presence or absence of a unique structure, other character concepts might require more careful consideration. In fossils, shape characters often contribute to an initial intuition about their placement, while they can be difficult to translate properly into discrete character states. Other characters might be omitted out of convenience or due to some uncertainty associated with the interpretation of the fossil. This was probably the cause of the unexpected placement of *Xanthopimpla messelensis*. This fossil showed a higher probability of belonging to Acaentinae than Pimplinae, and within the latter was rather associated with the sister genus *Lissopimpla* than with *Xanthopimpla* (Fig. 9c). Several characters that were used for the placement in the original description were not or not adequately represented in the morphological matrix: the enlarged claws (delimitation of this character was deemed too difficult and it was thus not included in the matrix), the potentially twisted mandible and short hypopygium (due to some uncertainty in the interpretation), the very narrow triangular areolet (as it is somewhat difficult to delimitate from other triangular areolets), and the slightly medially down-curved ovipositor (because it was not present in any other scored taxon). Similar shapes of the areolet and ovipositor occur in some extant *Xanthopimpla* species [62], and our failure to include them in the morphological matrix might have hindered the placement of the fossil. To achieve a more accurate estimate of the placement uncertainty in this fossil, a review of the character matrix would be warranted. When it comes to characters pertaining to shape and general ‘habitus’ of a taxon, morphometric approaches might improve current estimates.Continuous characters can in principle as easily be combined with discrete, morphological characters as the latter with molecular data [63], provided the respective models are implemented in the analysis software [60,64,65].

#### Insufficient taxon sampling

The estimate of the posterior probability of a taxon placement in a phylogeny can of course only take into account taxa that have in fact been sampled. If the closest relatives of a fossil are not included in a data matrix, then posterior probabilities are not actually conclusive, as has been the case for the fossil *Eclytus*? *lutatus* included here. As no extant representative of the genus *Eclytus* was sampled, the high probability of the fossil being associated with Orthocentrinae cannot be taken at face value, even though this alternative placement should be considered further in the future. Even if all potentially close relatives are included in an analysis, omission of other taxa might bias posterior probability, for instance if a certain character state is unique among the sampled taxa, but occurs in some not sampled groups as well. Not including these groups will imply a lower evolutionary rate of the character than is actually the case and thus overconfidence in the fossil placement. Adequate taxon representation is thus crucial for a meaningful morphological phylogenetic analysis.

#### Fossilization artefacts

If the fossilization process favours the preservation of ancestral character states in comparison with those that have evolved rather late in the history of a group, or if taphonomic loss of a character is erroneously interpreted as its absence, fossils might show more ancestral placements in a phylogeny than warranted, a phenomenon referred to as ‘stem-ward slippage’ [46]. While detailed studies of insect decay are missing, the fact that they lack biomineralized tissues means that they need exceptional conditions in order to fossilize, such as fine-grained lacustrine or shallow marine environments [6]. There is usually a pronounced difference in the preservation of different body parts, with the flat wings often exceptionally well preserved, including details of their venation, while the head with its compound eyes shows strong signs of decay; furthermore, legs and other appendages are often the first to dislocate from the body and are thus lost often for interpretation. However, it remains unclear whether these biases are aligned with the taxonomic significance of the character states, especially in ichneumonids. Characters used to diagnose ichneumonid subfamilies include characters from all over the body, including wing venation as well as carination on the body or structure of the ovipositor; it thus remains to be shown whether stem-ward slippage is an issue in this group, for instance causing the ancestral position of the poorly preserved *Dolichomitus*? *saxeus* (Fig. 4d).

#### Limited phylogenetic signal

Molecular phylogenies have all but replaced morphological phylogenies in the recent literature, of course unless fossils are included, and this transition has happened mostly because the phylogenetic signal in morphological data is believed to be relatively limited and prone to biases due to convergence and homoplasy [66,67,68]. The Ichneumonidae tree is no exception here: while most subfamilies have been recovered in our analysis, the backbone of the tree is very poorly resolved, which causes severe limitations for the interpretation of fossil placement. For instance, the higher-level groupings Ophioniformes, Ichneumoniformes, and Pimpliformes were not recovered here, which precludes us from even obtaining an estimate of the probabilities that a fossil belongs to a stem lineage of either of the three subfamily groups. This is for instance the case for *Eopimpla grandis* or *Mesornatus markovici*, both of which might be stem-lineage representatives of certain subfamilies or even of more ancestral lineages. As the phylogenetic uncertainty in this area of the tree is very high, no single branch that would potentially resolve the polytomy at the base of Ichneumonidae without Xoridinae obtained high attachment probabilities of the fossils.

In the case of ichneumonids, the limited phylogenetic signal in the morphological data is partly due to the high levels of homoplasy reported for the group (Gauld & Mound 1982). Many character states, especially those related to parasitoid life style and host choice, converged in distantly related groups (e.g., long ovipositor in parasitoids of wood boring insects; short and thick legs in parasitoids of concealed hosts, etc.). Another example is homoplasy due to similar size, which influences several character systems, leads to correlation among multiple characters in morphological matrices. For example, miniaturisation is often followed by a reduction of certain wing veins, changes in the shape of wing cells and pterostigma, and an overall stouter body. Such correlated morphological characters should in general be avoided, as much as possible, in phylogenetic analyses; but they can also be recognised as fast evolving and ‘down-weighted’ in Bayesian inference when enough information is present. *Eclytus lutatus* might be a case where a lack of information to infer homoplasy due to size (no data for the genus *Eclytus* or other small Tryphoninae genera) biased its placement in favour of Orthocentrinae, a subfamily of small to very small ichneumonids.

A potential solution when confronted with limited phylogenetic signal in a morphological matrix would be to perform a combined analysis with molecular data in order to get a better-resolved consensus tree on which fossil placement could then be plotted; this is certainly a valid alternative approach. However, we decided to use morphology only, and even only adult, external morphological characters which are potentially visible in fossils, because we this way obtained a consensus tree which in itself shows the limits of achievable fossil placement certainty: it demonstrates the limitations of the phylogenetic signal in the morphological dataset even for extant taxa, for which many more characters can be scored than for fossils. Interestingly, however, our morphological tree is not in fact that much less resolved than a molecular tree resulting from a recent 93-gene study, with the exception of the three afore-mentioned subfamily groups [45]. The signal in the morphological data is thus not even that much weaker, and the addition of larval characters would likely even improve resolution along the backbone of the tree [69].

#### Model misspecification

While testing the ‘unordered’, ‘ordered’ and ‘full-state’ Mk models, we were not able to include the full range of models that have been developed for morphology, as this would have been beyond the scope of this study [12,14,15]. However, we would like to point out a potential artefact caused in our data by model misspecification that pertains to the stationarity assumption [13] and concerns the two species of *Crusopimpla* (Figs 4b, c). This genus was erected as a stem-lineage representative of the subfamily Pimplinae, with the presence of several carinae on the propodeum as a defining characteristic. None of the extant pimplines has complete carination (even though it is rather extensive in some Theronini), which might explain why such a placement was never chosen. However, propodeal carination might evolve in a very non-stationary manner in Ichneumonidae: the earliest branching subfamily, Xoridinae, has complete carination of the propodeum, and the character distribution on the phylogeny [28] suggests that this carination has progressively been reduced in parallel in several of the more derived ichneumonid subfamilies, including Pimplinae. The fact that different sets of propodeal carinae are reduced in different tribes of Pimplinae provides further support for the hypothesis that its ancestor might have had a more complete carination. If this is indeed the case, it would require the use of a directional model of character evolution, such as the one used to study the reduction in hymenopteran wing venation and muscles [13]. However, this model is not currently available for multistate data.

### Future directions

Model-based phylogenetics already represents a powerful tool for assessing the relationships among fossil and extant taxa, and the future developments of models of morphological evolution are likely to further improve these estimates. Besides relaxing the assumptions of equal rates among characters [43], symmetry of transition rates [15], and process stationarity [13], morphological data might also evolve with higher differences in effective branch lengths among characters than molecular data [57], even though this remains to be demonstrated. A no-common-mechanism model for morphology [70] would quickly be beyond the scope of any stochastic analysis approach and lead to a strong discrepancy between the number of parameters in the model and available data points from which to estimate them. However, if morphological data can be partitioned according to biological characteristics such as functional units, using separate sets of branch lengths for each partition might still be tractable. An equivalent approach has been taken in the context of a calibrated phylogeny of mammals, where different morphological partitions were allowed to evolve according to a separate relaxed-clock model [71].

Another natural extension of our approach would include the ages of fossils to improve their placement, either in the context of morphology-only tip dating [19,20] or combined with molecular data in total-evidence dating [23,72]. This might have a considerable impact; for instance, knowing that a fossil is older than the inferred splits in the crown group would enforce its placement as a stem-lineage representative. However, including age information might interact with the morphology model to produce artefacts, especially if a character evolves far from stationarity on the time scale covered by the phylogeny [13]. Carful model testing would expose such artefacts and improve our understanding of character evolution. In any case, using the context of a calibrated phylogeny to further test the placement of ichneumonid fossils would represent a natural next step and lead to a better understanding of the evolutionary history of this highly species-rich taxon.

## Acknowledgements

We thank Gavin Broad (NHM London) for extensive discussion of morphological character conceptions. Emmanuel Paradis (Université de Montpellier) provided plotting functions of trees in R, which we modified to produce RoguePlots. For access to the fossils covered in this article, we are grateful to Marsh Finnegan and Alan M. Rulis (Smithsonian National Museum of Natural History, Washington, USA), Talia Karim and David Zelagin (Museum of Natural History, University of Colorado, Boulder, USA), Ricardo Pérez-de la Fuente (Museum of Comparative Zoology, Harvard University, USA), Dmitry Kopylov (Borissiak Paleontological Institute, Russian Academy of Sciences, Moscow, Russia) and Sonja Wedmann (Forschungsstation Grube Messel, Senckenberg, Germany). Extant specimens were made available by the Swedish Malaise Trap Project (www.stationlinne.se, Sweden), Eric Chapman (University of Kentucky, U.S.A.), Masato Ito (Kobe University, Japan), Ilari Sääksjärvi (University of Turku, Finland), and Gavin Broad (NHM London). This study was funded by the Swiss National Science Foundation (grant PZ00P3_154791 to SK).

## Supplementary material

Supplementary File S1. Morphological matrix in NEXUS format.

Supplementary File S2. Full list of all coded extant and fossil taxa. Extant specimens were labeled with a project number (Ichn_#XXXX) and are deposited at the specified collections. Collection abbreviations: KYU (Department of Entomology, University of Kentucky, USA), MCZ (Muzeum of Comparative Zoology Harvard), MNHN (Museum National d’Histoire Naturelle), NHM (Natural History Museum London), NHRM (Naturhistoriska Rijkmuseet Stockholm), NMB (Naturhistorisches Museum Basel, Switzerland), NMBE (Naturhistorisches Museum Bern, Switzerland), NMNH (National Museum of Natural History Smithsonian), PIN (Paleontological Institute of Russian Academy of Science), SMF (Senckenberg Forschungsinstitut und Naturmuseum), UCM (University of Colorado Museum of Natural History), WC (Waite Insect collection, University of Adelaide, Australia), ZMUT (Zoological Museum of the University of Turku, Finland). For the fossils collections, we provide the collection’s individual numbers. The gender of the coded specimens was indicated by (f) for female and (m) for male.

Supplementary File S3. Description of morphological characters. Illustrated list of the morphological characters and their states used in this study.

Supplementary File S4. R functions to create RoguePlots and to prepare morphological matrices that include polymorphisms for analysis.

Supplementary File S5. Complete RoguePlots of the individual fossils.

